# The tendon interfascicular basement membrane provides a vascular niche for CD146^+^ pericyte cell subpopulations

**DOI:** 10.1101/2022.10.14.512258

**Authors:** Neil Marr, Danae E. Zamboulis, Dirk Werling, Alessandro A. Felder, Jayesh Dudhia, Andrew A. Pitsillides, Chavaunne T. Thorpe

## Abstract

The interfascicular matrix (IFM) is critical to the mechanical adaptations and response to load in energy-storing tendons, such as the human Achilles and equine superficial digital flexor tendon (SDFT). We hypothesized that the IFM is a tendon progenitor cell niche housing an exclusive cell subpopulation. Immunolabelling of equine SDFT was used to identify the IFM niche, localising expression patterns of CD31 (endothelial cells), CD146 (IFM cells) and LAMA4 (IFM basement membrane marker). Magnetic-activated cell sorting was employed to isolate and compare *in vitro* properties of CD146^+^ and CD146^-^ subpopulations. CD146 demarcated an exclusive interfascicular cell subpopulation that resides in proximity to a basal lamina which forms interconnected vascular networks. Isolated CD146^+^ cells exhibited limited mineralization (osteogenesis) and lipid production (adipogenesis). This study demonstrates that the IFM is a unique tendon cell niche, containing a vascular-rich network of basement membrane, CD31^+^ endothelial cells and CD146^+^ cell populations that are likely essential to tendon structure- and/or function. Interfascicular CD146^+^ subpopulations did not exhibit stem cell-like phenotypes and are more likely to represent a pericyte lineage. Previous work has shown that tendon CD146 cells migrate to sites of injury, therefore mobilisation of endogenous tendon IFM cell populations may promote intrinsic repair.

## 1. Introduction

Tendons are fundamental components of the musculoskeletal system, acting as connections between muscle and bone. The predominant function of tendon is to transfer the forces exerted by skeletal muscle contractions to bone, positioning the limb for locomotion [1,2]. However, specialised energy-storing tendons, such as the equine superficial digital flexor tendon (SDFT) and human Achilles tendon, enhance the functional adaptation of tendon by lowering the energetic cost of locomotion through their mechanical properties, such as greater extensibility, elasticity and fatigue resistance [3-5]. Much like skeletal muscle, these specialised mechanical properties of energy-storing tendons are provided by their hierarchical structure of subunits predominantly composed of type I collagen, forming fascicles which are surrounded and bound by a non-collagenous interfascicular matrix (IFM) which governs the high-strain behaviour of energy-storing SDFT by facilitating sliding between fascicles [6]. While the mechanical role of IFM in the function of energy storing tendons is well defined, less is known regarding its biological role in developing, adult and ageing tendon, particularly regarding the identity and function of IFM localised cell populations and their niche. Histological analyses of tendon have revealed regional morphological differences in cell populations, with rounder cells within the IFM, present in greater number compared to those within the fascicles which are highly aligned with the long axis of the tendon [7,8]. In addition, seminal studies have alluded to an endogenous tendon stem/progenitor cell (TSPC) population and niche, both of which remain largely undefined but have been speculated to reside within the IFM [9-11]. In other tissues, the stem cell niche is maintained by mechanically unique microenvironments, similar to the high shear environment within the IFM, which may therefore be the location of the tendon stem/progenitor cell niche [12-14].

Tendon development is driven by stem/progenitor cell populations which express Mohawk homeobox (MKX) and Scleraxis (SCX) transcription factors [15-17], however their intracellular localisation impedes cell sorting techniques required for *in vitro* study of stem/progenitor cell populations. In adult tissues, cell surface markers, such as CD44 and CD90 (THY1), are members of a number of canonical marker panels routinely used in the characterisation and isolation of specific stromal/stem cell populations [18,19]. Recent studies have also reported resident CD146 populations in tendon [20,21]. CD146 or melanoma adhesion molecule (MCAM; MUC18; Gicerin; OMIM:155735) is a transmembrane glycoprotein belonging to the IgG superfamily of cell adhesion molecules [22]. Originally characterised as a marker of tumour progression and metastasis, CD146 has since been reported as a marker of 4 (LAMA4) [30]. However, few endothelial cell lineages, both haematopoietic and mesenchymal stem cell lineages, as well as synovial fibroblasts and periosteal cells [23-29]. Our laboratory has recently reported that these CD146 subpopulations are present within the IFM of the rat Achilles and recruited to injury sites from their IFM niche via the CD146 ligand Laminin α studies have attempted to comprehensively characterise CD146 tendon cells and their *in vivo* cell niche composition.

In this study, we tested the hypothesis that the IFM is a tendon progenitor cell niche housing an exclusive cell subpopulation. We report novel markers of interfascicular cells and basement membrane, and identify CD146 as an optimal marker for use in IFM cell sorting procedures. We also demonstrate that the lineage potential and clonogenicity of interfascicular CD146 cells is limited, which may be indicative of a differentiated vascular population in contrast to resident tendon stem/progenitor cells.

## 2. Materials and Methods

### 2.1. Ethical statement

The collection of animal tissues was approved by the Royal Veterinary College Ethics and Welfare Committee (URN-2016-1627b). All tissues were sourced from horses euthanised for reasons other than tendon injury at a commercial equine abattoir.

### 2.2 Tissue acquisition

Superficial digital flexor tendons (SDFT) were harvested from forelimbs taken from young, skeletally mature horses (age=3-8 years, n=5, exercise history unknown). Prior to isolation, the forelimbs were clipped to remove hair and the skin sterilised by several applications of 4% chlorhexidine (HiBiScrub®; Mölnlycke Health Care). Portions of mid-metacarpal SDFT (6-10 cm) were dissected free of the limb and stored immediately in standard growth medium consisting of pyruvate and low glucose Dulbecco’s modified eagle medium (DMEM) supplemented with 1% (v/v) penicillin/streptomycin and 10% (v/v) qualified, heat-inactivated foetal bovine serum (FBS) until tissue processing (all from Gibco™). Excised tendons presenting with previously reported definitions of macroscopic evidence of injury were excluded from all experiments [31,32]. Dissections and subsequent cell processing were completed within 24 h of euthanasia.

### 2.3 Cryosectioning

SDFT frozen sections were prepared as previously described [33]. Tissues were briefly washed in Dulbecco’s phosphate-buffered saline (calcium and magnesium free, embedded with optimal cutting temperature compound (OCT; Cell Path, Newtown, UK) embedding matrix and snap-frozen in pre-cooled hexane on dry ice. Serial longitudinal sections of 6-20 μm thickness were prepared using a cryostat microtome (OTF5000, Bright Instruments) equipped with MX35 Premier Disposable Low-Profile Microtome Blades (3052835, Fisher Scientific). Tissue sections were mounted on SuperFrost™ Plus Slides (10149870, Fisher Scientific), air-dried at room temperature (RT) for a maximum of 2 h and stored at −80 °C.

### 2.4 Periodic acid-Schiff staining

Periodic acid-Schiff (PAS) staining was used to detect mucins and basement membrane proteins. Staining was performed using an Alcian Blue (pH 2.5)/PAS staining kit according to manufacturer guidelines (Atomic Scientific). SDFT cryosections (20 μm) were thawed and fixed with 4% PFA/10% NBF for 10 mins at RT. Slides were rinsed thoroughly with distilled water, stained with 1% Alcian blue in 3% acetic acid (pH 2.5) for 10 mins, and washed thoroughly in distilled water. Slides were treated with 1% periodic acid solution for 10 minutes at RT, washed with distilled water, then treated with Schiff reagent (Feulgen) for 10 minutes at RT. Sections were then washed under running tap water until sections presented a magenta colour macroscopically. Sections were then counter-stained with haematoxylin, dehydrated and cleared using an automated slide stainer (Varistain^™^ Gemini ES), and mounted with glass coverslips using DPX mountant. Slides were cured at RT overnight and imaged using brightfield microscopy (DM4000B upright microscope) in Leica Application Suite software version 2.6 (Leica Microsystems).

### 2.5 Network-based predictions of CD146 interactions

Proteins of interest for immunolabelling were selected based on their expression by tendon progenitor cell subpopulations in previous reports [21] and their predicted interactions in *Equus caballus* (NCBI taxid: 9796) using STRING (version 10.5) network-based predictions for CD146 and LAMA4 [20,34].

### 2.6 Immunolabelling

SDFT cryosections were thawed and fixed with acetone (pre-cooled at −20 °C) for 10 mins, washed three times for 5 mins at RT with tris-buffered saline (TBS), incubated in ‘blocking’ buffer (TBS supplemented with 1% (w/v) bovine serum albumin (Scientific Laboratory Supplies), 5% (v/v) goat serum (Sigma), and 5% (v/v) horse serum (Sigma)) for 2 hours. Horse serum was used to saturate Fc receptors on the surface of cells within the tissue.

Sections were incubated with primary antibodies overnight at 4 °C (details regarding primary and secondary antibodies are provided in **Table S1**). For negative controls, sections were treated with blocking buffer only. For isotype controls, sections were treated with mouse and rabbit IgG isotype-matched controls diluted in blocking buffer at identical concentration to primary antibodies used.

For fluorescent detection (10 μm sections), secondary antibodies diluted in blocking buffer were applied to sections and incubated for 1 h at RT under dark conditions. Sections were washed with three times with TBS for 5 mins, and mounted with glass coverslips using ProLong^™^ Diamond antifade mountant with 4’,6-diamidino-2-phenylindole (DAPI) as a nuclei counterstain. Slides were cured for 24 h at RT under dark conditions, prior to imaging. Negative and isotype matched control images for fluorescent labelling are provided in **Figure S1**.

Immunohistochemical labelling (6 μm sections) was performed in a similar manner to fluorescent detection, using an EnVision®^+^ Dual Link System-HRP DAB^+^ system (Dako), with the inclusion of an of EnVision dual endogenous enzyme block for 15 mins at RT under dark conditions prior to treatment with blocking buffer, and wash steps were performed using 0.05% (v/v) TBS-TWEEN20. For immunohistochemical detection, sections were incubated in EnVision peroxidase labelled polymer (conjugated to goat anti-mouse and goat anti-rabbit immunoglobulins) for 30 mins at RT. Sections were then washed three times and incubated with EnVision DAB^+^ substrate buffer-3,3’-diaminobenzidine (DAB) chromogen solution for 3 mins, rinsed three times with deionised water (diH2O), counter-stained using haematoxylin according to Delafield, dehydrated and cleared using standard procedures on a Varistain^™^ Gemini ES automated slide stainer, then finally mounted with glass coverslips using DPX mountant. Slides were cured at RT overnight and imaged using brightfield microscopy (DM4000B upright microscope) in Leica Application Suite software version 2.6 (Leica Microsystems). Negative control images for immunohistochemistry are provided in **Figure S2**.

### 2.7 Fluorescent labelling analyses

To distinguish between regions of IFM and fascicular matrix (FM), boundaries between both phases were determined by light refraction in phase contrast images, as well as gross identification by nuclei number and cell morphology (**Figure S3**).

For quantification, all settings remained constant between samples including exposure, pixel size, z-step size, and laser settings with all images taken in one single session. For each sample, two distinct areas were imaged in two separate serial tissue sections (2× sections per horse donor, n=5).

Confocal images are presented as maximum intensity projections from z-stacks containing image slices at a resolution of 512 × 512 × 40 pixels (227.9 × 227.9 × 13.09 μm; 0.34 μm z-step size) to fully capture tissue depths. Image processing and analysis was performed using Fiji/ImageJ software [35]. For IFM measurements, an area fraction (%) of positively stained pixels were recorded in 8-bit binary images (black = negative, white = positive) to measure expression of markers of interest. To generate binary images for each marker, a background correction was performed to remove noise, followed by a median filter and threshold (Triangle for CD146/MKX = 555 nm, Huang for CD44/CD90 = 633 nm). The lookup table (LUT) of colour channels within images was changed for visualisation purposes.

### 2.8 3D immunolabelling

3D immunolabelling of SDFT segments was performed as previously described [36]. All steps were performed with orbital agitation. SDFT segments (5□mm□×□5□mm□×□2□mm) were washed twice for 12 h with TBS at RT, and permeabilised sequentially in 50% (v/v) methanol:TBS, 80% (v/v) methanol:diH2O, and 100% methanol for 2 h at 4 °C. Samples were washed sequentially for 40 minutes at 4 °C with 20% (v/v) DMSO:methanol, 80% (v/v) methanol:diH2O, 50% (v/v) methanol:TBS, TBS, and TBS supplemented with 0.2% (v/v) Triton X-100. Prior to blocking, samples were incubated with a pre-blocking penetration buffer containing 0.2% TBS-TX100, 0.3 M glycine, and 20% DMSO for 6 h at 37 °C. Equine SDFT segments were blocked for 80 h at 37 °C in 0.2% TBS-TX100 supplemented with 6% (v/v) goat and 6% (v/v) donkey serum and 10% (v/v) DMSO. Primary antibody incubations for CD146 (1:100) were performed at 37 °C for 80 h in wash buffer (TBS supplemented with 0.2% (v/v) TWEEN20), 3% (v/v) goat serum, 3% (v/v) donkey serum, and 5% (v/v) DMSO. Segments were washed 3× 2 h with wash buffer, incubated with secondary antibodies (1:250, goat anti-rabbit Alexa Fluor® 594) for 36 h at 37 °C, washed 5× 5 mins with wash buffer, and counterstained overnight with DAPI (1:2000) diluted in wash buffer. Segments were dehydrated with increasing concentration of methanol, and tissue cleared with immersion in Visikol® HISTO™-1 for 36 h, followed by immersion in HISTO™-2 for at least 36 h at RT. Samples were stored in HISTO™-2 at 4 °C prior to confocal imaging.

Confocal imaging of regions (approx. 1□mm□×□1□mm□×□0.2□mm) within each sample was performed using a Leica TCS SP8 laser scanning confocal microscope with a motorised stage. Our previous studies have established antibody penetration of at least 0.2 mm in SDFT segments [37]. Images were captured using lasers emitting light at 405 (blue channel; DAPI) and 561 (red channel; Alexa Fluor 594) nm with laser power <□10% and scanning speed = 600□Hz with a HC PL FLUOTAR 10x/0.32 dry objective lens, resolution = 1024□× □1024 px, pinhole size = 1 Airy unit, frame average = 1, line average = 8, and electronic zoom = 0.75. 3D renderings were captured in Leica LAS X software (version 3.5.5) within the 3D module.

### 2.9. Primary tendon cell culture

SDFTs collected under sterile conditions were placed in Petri dishes containing Gibco™ Dulbecco’s PBS (without phenol red, calcium and magnesium) supplemented with 1% (v/v) antibiotic-antimycotic solution. Surrounding peritenon was removed to isolate the tendon core (6 g), which was diced into approximately 4 mm^3^ pieces, rinsed with DPBS, and digested with 1 mg/mL pronase E (39052, VWR) per 1 g tissue for 6-8 h at 37 °C and 5% CO_2_ under constant agitation. Following pronase digestion, tissue was digested for a further 24 h with 0.5 mg/mL collagenase type IV (CLS-4, Lorne Laboratories) and 1 mg/mL dispase II (17105041, Invitrogen) at 37 °C and 5% CO_2_ with constant agitation [38].

### 2.10. Magnet-activated cell sorting (MACS) of CD146 cells

Previous studies have shown that >50% expression of cell membrane proteins can be restored post-digestion by 24 h *in vitro* culture [39]. Hence, to enhance antigen recovery, freshly digested tendon-derived cells (TDCs) were cultured overnight to maximise CD146 cell isolations. Following this recovery phase, adherent cells were dissociated at 37 °C for 10 mins using Accutase® solution according to manufacturer’s guidelines. Cells remaining in suspension (i.e. non-adherent populations) were also collected alongside dissociated cells (adherent populations). Cell isolates were washed by resuspension in fresh growth medium and centrifuged at 300 × g for 10-20 mins depending on pellet formation. Cell pellets (passage 1; p1) were resuspended in growth medium and separated into single-cell suspensions (SCSs) by passing through a 70 μm cell strainer. SCSs were resuspended in freshly prepared, ice-cold MACS buffer containing sterile-filtered FACSFlow™ (342003, BD Biosciences) supplemented with 1% (w/v) BSA. SCSs were centrifuged for 10 mins at 300 × g, resuspended in MACS buffer, and both cell viability and numbers determined by trypan blue (T8154, Sigma-Aldrich) and a haemocytometer. Suspensions with <90% viability were discarded. SCSs were incubated with anti-CD146 antibodies (ab75769, Abcam, Cambridge, UK) at a concentration of 1 μg/mL for 30 mins at 4 °C on ice.

Following primary antibody incubation, SCSs were washed three times by centrifugation at 300 × g, resuspended in MACS buffer, and incubated with anti-rabbit IgG micro-beads (130-048-602, Miltenyi biotec) diluted in MACS buffer for 15 mins at 4 °C. SCSs were washed three times by centrifugation at 300 × g and resuspended in MACS buffer.

MidiMACS™ LS columns (130-042-401, Miltenyi biotec) were mounted to a MidiMACS™ Separator and multistand (130-042-301, Miltenyi biotec) and washed with MACS buffer according to manufacturer guidelines. MACS-ready SCSs were passed through MidiMACS™ columns and washed with MACS buffer twice. All wash elutions containing negatively selected cells (i.e. CD146^-^ TDCs) were collected on ice until processing of sub-cultures. Following negative cell depletion, CD146^+^ cells were collected by removing the MACS column from the MACS magnet and eluting the column with MACS buffer and a plunger.

All sub-cultures were maintained until a maximum of three passages (p3) to limit phenotypic drift. For downstream assays, cells were dissociated using Accutase® solution (A6964, Sigma-Aldrich).

### 2.11. Flow cytometry

For direct flow cytometry, 0.1-0.2×10^6^ cells were resuspended in DPBS. For CD146^+^ cells, lower concentrations were used according to yields following MACS isolation. All tubes were stored on ice immediately prior to and during flow cytometry. Cell suspensions (50 μL) were incubated with a phycoerythrin (PE)-conjugated variant of the EPR3208 anti-CD146 antibody (1:100, ab209298, Abcam, Cambridge, UK) on ice for 30 mins, washed with DPBS and spun at 400 × g. Supernatant was removed, and pellets resuspended in 500 μL DPBS for immediate flow cytometry analyses.

All flow cytometry acquisition was performed using an air-cooled 3-laser BD FACSCanto II™ flow cytometer (BD Biosciences) equipped with BD FACSDiva (version 8.0.1, BD Biosciences). Acquisition equipment and software were calibrated daily or immediately prior to acquisition using BD FACSDiva™ CS&T Research Beads (BD Biosciences). Data analyses was performed in FlowJo software (version 10, FlowJo LLC). Unstained controls (fluorescence minus one control) were used to gate and discriminate positively and negatively labelled populations. The percentage of positive cells gated in unstained samples (i.e. autofluorescent cells) was subtracted from stained samples (i.e. experimental cells) to give an overall percentage of immunoreactivity. All experiments recorded a minimum of 10,000 total events (i.e. cells).

### 2.12. Immunocytochemistry

For detection of CD146 in unsorted TDCs, 0.1-0.2×10^6^ cells were seeded on sterile 16 mm borosilicate glass circle coverslips coated with poly-L-lysine solution (0.01%, sterile-filtered, P4832, Sigma-Aldrich) until 70-80% confluence. To detect CD146 within MACS-enriched CD146^+^ cells, immunocytochemistry was performed directly on cells (0.1-0.2×10^6^ seeding density) in non-coated culture vessels at 70-80% confluence. Cells were washed 3 times with DPBS, fixed with pre-chilled (−20 °C) acetone:methanol (1:1) for 20 mins on ice, then washed three times with DPBS. Cells were blocked for 1 h with blocking buffer as described above. Cells were incubated overnight with primary antibodies overnight at 4 °C as described above, washed three times with DPBS, incubated for 1 h with secondary antibodies (1:500, goat anti-rabbit Alexa Fluor® 488 and goat anti-mouse Alexa Fluor® 594).

For direct CD146 labelling in MACS-sorted populations, cells were incubated overnight at 4 °C with phycoerythrin (PE)-conjugated anti-CD146 antibodies (1:100, ab209298, Abcam, Cambridge, UK). Cells were washed three times with DPBS, labelled with DAPI (1 μg/mL) for 2 mins, washed three times with DPBS and mounted using Prolong™ Diamond, cured at RT under dark conditions for 24 h before storing at 4 °C until imaging. Fluorescent imaging of TDCs was performed using a Leica SP5 (40× HCX PL FLUOTAR PH2 NA=0.75 objective). For CD146^+^ cells, imaging was performed on a DMIRB inverted microscope (Leica Microsystems, Wetzlar, Germany; 40× N PLAN L corr PH2 NA=0.55 objective).

### 2.13. Clonogenic assay

Bone marrow-derived mesenchymal stromal cells (MSCs) isolated as described previously were kindly provided by Dr Giulia Sivelli [10]. MSCs, unsorted TDCs, CD146^-^ cells and CD146^+^ cells were seeded in 6-well plates at a density of 100 cells cm^-3^ (approx. 900 cells) and cultured for 7 d. At termination of cultures, cells were washed 3× with DPBS, fixed with 2.5% glutaraldehyde for 10 mins, then washed 3× DBPS (all steps at RT). Cells were stained with 0.1% (v/v) crystal violet for 30 mins at RT. Cells were washed 3x with DBPS and left to air dry at RT. Images were acquired using a flat-bed scanner (Epson Perfection 4990, Epson) at a resolution of 800 dpi.

### 2.14. Adipogenesis assay

MSCs, unsorted TDCs, CD146^-^ cells and CD146^+^ cells were seeded into 12-well plates at a density of 0.4×10^5^ cells per well and cultured for 48 h until adherence in standard growth medium. To induce adipogenesis, standard growth media was removed, and cells were cultured with StemPro® Adipogenesis differentiation media for a further 14 d. Cells were fed induction media every 72 h. Upon termination of culture, monolayers were washed once with DPBS before fixation with 4% PFA/10% NBF for 30 mins at RT.

To assess intracellular lipid vesicles produced by adipogenic conditions, cells were stained with Oil Red O. Fixed monolayers were rinsed once with distilled water then washed with 60% isopropanol for 5 mins at RT. Monolayers were stained for 15 mins at RT with a 3:2 working solution of 3-parts 0.3% (w/v) Oil Red O diluted in isopropanol and 1-part distilled water. Cells were washed repeatedly with distilled water until rinsed clear of precipitating Oil Red O, then counterstained with Harris haematoxylin for 1 min at RT.

Imaging was performed on an Axiovert 135TV inverted microscope (Zeiss) using Image Pro Insight version 9.1.4 (Media Cybernetics).

### 2.15. Osteogenesis assay

Following dissociation, MSCs, unsorted TDCs, CD146^+^ and CD146^-^ cells were seeded into 12-well plates a density of 0.1×10^6^ cells per well with osteogenic media containing 2 mM sodium phosphate dibasic (DiP) or standard growth medium as a control with each condition supplemented with 50 μg/mL ascorbic acid to promote collagen synthesis [40,41]. DiP (free phosphate donor) is essential for bone/mineralised extracellular matrix metabolism during osteogenesis [42]. Monolayers were fed with fresh half-media changes corresponding to each condition every 72 h.

Cell cultures were terminated after 21 days to assess mineralisation with Alizarin Red S staining [43]. Monolayers were rinsed once with DPBS then fixed for 10 mins at RT with 2.5% (v/v) glutaraldehyde. Fixed cells were rinsed once with DBPS then three times with 70% ethanol and air-dried at RT overnight. Dried monolayers were subsequently stained with 1% (w/v) Alizarin Red S in diH2O for 5 mins at RT, then washed three times with 50% ethanol and left to air-dry overnight. Imaging was performed as described above.

### 2.16. Statistical analyses

Statistical analyses and graphs were produced using GraphPad Prism (version 9.1). Normality tests were performed according to Shapiro-Wilk tests (α=0.05). All datasets passed normality tests and were analysed using unpaired two-tailed t-test (significance set to P <0.05). Graphs were plotted as mean (μ) ± standard deviation (SD).

## 3. Results

### 3.1 CD146 is a marker of interfascicular cell populations

PAS staining demonstrated that the IFM contains mucin-rich basement membrane. Using both CD146 and the IFM basement membrane marker LAMA4 in STRING predictions identified several potential interfascicular cell surface markers including CD44, CD90 (THY1) and CD133 (PROM1), as well as a broader network of interfascicular niche and basement membrane components, including dystroglycan 1 (DAG1), integrin subunit β1 (ITGB1) and fibronectin 1 (FN1) amongst other potential proteins of interest (Fig. 1A-B).

**Figure 1.**
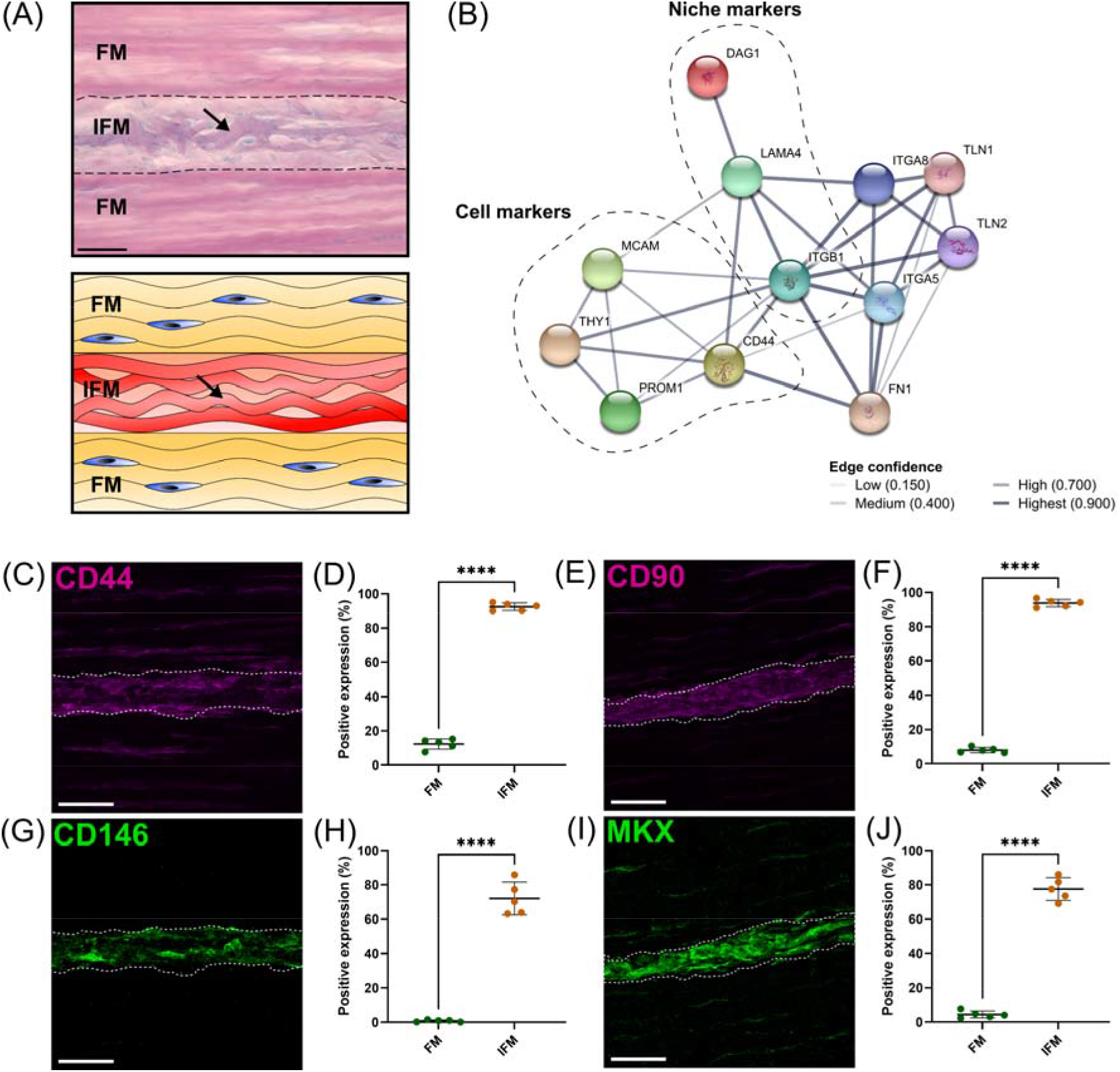
Analyses of regional differences in tendon cell marker expression demonstrated that CD146 is exclusively expressed by interfascicular cells within an interfascicular niche. (A) PAS-staining and schematic of SDFT sections highlighted mucin-rich basement membrane (arrow; purple) of vasculature (schematic; red) within the interfascicular matrix (IFM). Nuclei = blue. Scale bar = 50 μm. (B) STRING-predicted protein-protein interactions revealed potential targets for novel tendon cell populations using validated interfascicular niche markers CD146 and LAMA4. Interactions based on CD146 (MCAM) and LAMA4 demonstrated a protein neighborhood consisting of cell markers CD44, CD90 (THY1), CD133 (PROM1), as well as cell niche components such as ITGB1, DAG1 and FN1. (C-J) Image analyses comparing the positive labelling (area fraction; %) of longitudinal SDFT sections immunolabelled with CD44 (C,D), CD90 (E,F), CD146 (G,H) and MKX (I,J) in both fascicular matrix (FM) and IFM regions. The IFM is outlined by dotted lines. Scale bar = 50 μm. Biological replicates (n) = 5 per tendon region. Technical replicates = 3-4 per individual sample. Graphs were plotted as mean (μ) ± SD. Statistical significance: **** (P ≤ 0.0001).

To validate these proposed interfascicular cell markers, fluorescent labelling of CD44, CD90, CD146 and MKX was quantified in both fascicular and interfascicular regions (Fig. 1C-J). All markers were enriched within IFM (72-94% positive expression) and had significantly less expression within fascicles, with fascicular CD146 expression less than 1% compared to CD44, CD90 and MKX which were between 4-15%.

Using 3D imaging of SDFT labelled with CD146, we identified an interfascicular network of vascular structures within which CD146 cells were localised (Fig.2A). The colocalisation of CD146 with CD31 (endothelial marker) or LAMA4 (basement membrane) (fig 2 B-C) confirmed that the structures were vascular.

**Figure 2.**
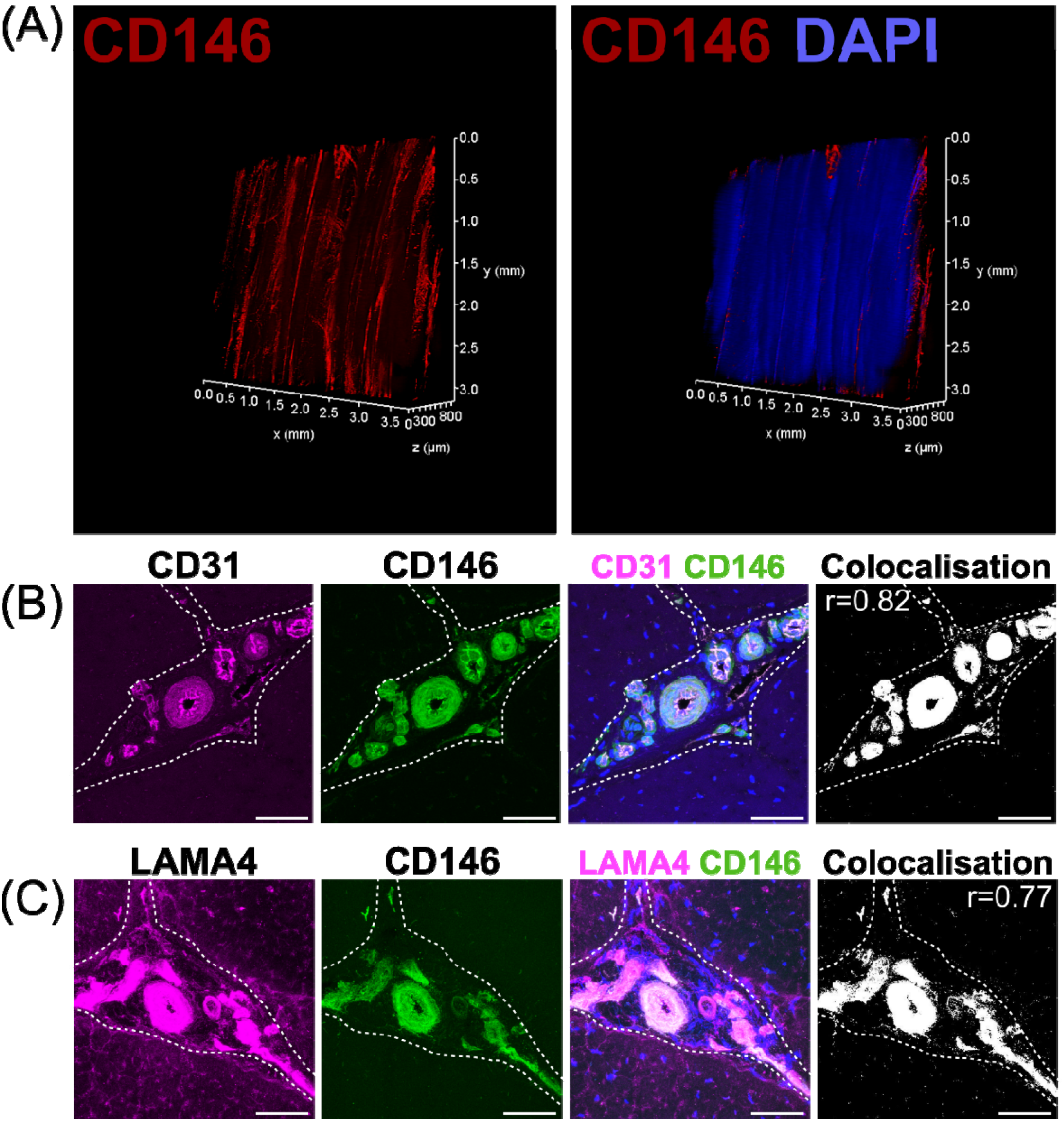
CD146 cell populations demarcated a vascular network indicative of a vascular cell niche within the interfascicular matrix. 3D imaging of CD146 (A) confirmed that the IFM was enriched with CD146 cell populations as part of an interconnected vessel network. Images of labelled transverse SDFT sections demonstrating colocalisation of CD31, CD146, and LAMA4 (B,C), indicating that CD146 represents a marker of interfascicular vascular/endothelial cell populations resident within laminin-rich vessels. Nuclei = DAPI (blue). IFM is demarcated by dashed lines in transverse images. Scale bar = 50 μm.

### 3.2 Tendon interfascicular matrix is enriched in an endothelial basement membrane

To identify and characterise the major components of interfascicular basement membrane, we performed immunolabelling of basement membrane proteins, including full-length laminin (pan-laminin), type IV collagen, and Perlecan (Fig. 3A-C), all of which localised to the vasculature within IFM. Further labelling with endothelial markers endomucin (EMCN) and von willebrand factor (VWF) demonstrated abundant expression within the IFM (Fig. 3D-E). STRING predictions in *Equus caballus* identified canonical basement membrane components integrin β1 (ITGB1) and dystroglycan 1 (DAG1), as part of the CD146-LAMA4 interaction network. Hence, we performed labelling of α-dystroglycan (IIH6) and ITGB1 (Fig. 3F&H), in addition to network-predicted cell surface marker CD133 (Fig. 3G) with all three labelled abundantly within the IFM. As LAMA4/LAMA5 ratios are critical for basement membrane integrity [44], we also demonstrated labelling for laminin α5 (LAMA5) within the IFM (Fig. 3I). We also examined other reported angiogenic mediators, Netrin-1 (NTN1); a reported ligand of CD146, and Neuropilin-1 (NRP1), both of which also localised to the IFM (Fig. 3J-K).

**Figure 3.**
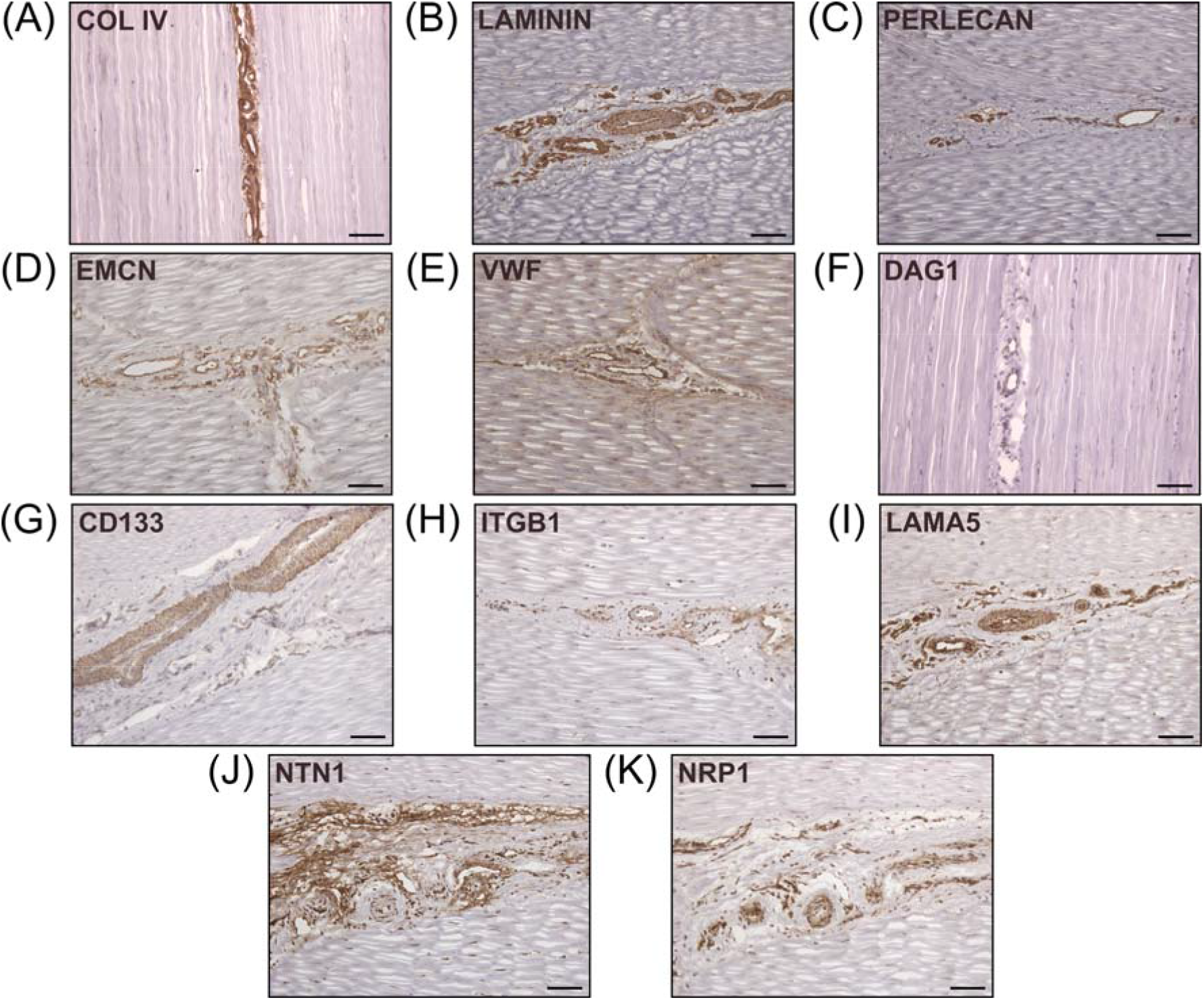
Canonical and network-predicted vascular basement membrane components were enriched within the interfascicular matrix. Experimental validation confirmed interfascicular expression of Type IV collagen (A), full-length laminin (B), Perlecan (C), as well as vascular markers EMCN (D) and VWF (E). Immunolabelling validation also confirmed enrichment within interfascicular vasculature with network-predicted markers DAG1 (IIH6; F), CD133 (G), IGTB1 (H), laminin isoform LAMA5 (I), and angiogenic mediators NTN1 (J) and NRP1 (K). Scale bar = 75 μm.

### 3.3 Interfascicular CD146^+^ cells are a rare subpoplation requiring enrichment for in vitro isolation

Upon isolation from the SDFT, *in vitro* labelling of cell surface markers demonstrated that the majority of TDCs exhibited abundant CD44 and CD90 labelling and limited CD146 expression (Fig. 4A-C). Reported TSPC marker Nestin (NES) was not detected in these *in vitro* cultures.

**Figure 4.**
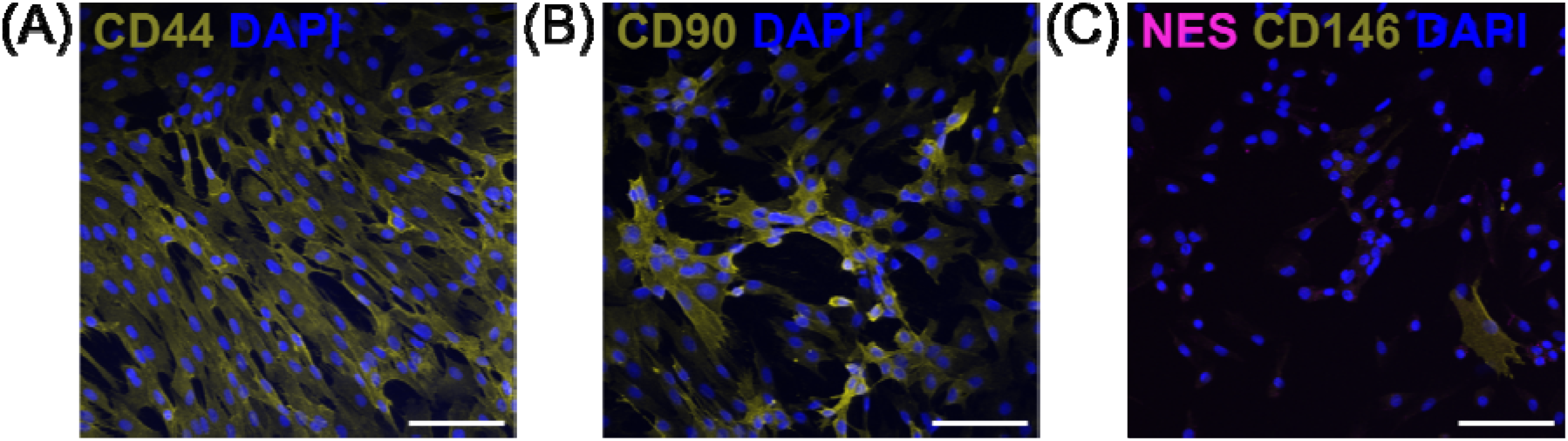
Immunocytochemical labelling confirms CD146^+^ cells are a rare subpopulation when expanded *in vitro*. Tendon cell-surface markers CD44 (A), CD90 (B) and CD146 (C) demonstrate that CD146 cells are a rare subpopulation amongst cultured TDCs when compared to cells expressing CD44 or CD90. No NES was detected in TDCs (passage 1). n=3 (biological replicates). Scale bar = 100 μm.

To study CD146 cells *in vitro*, we developed a MACS procedure for the enrichment of CD146 cells. A single application of MACS enrichment was able to yield CD146 cells with purity of approximately 64% as determined by flow cytometry (Fig. 5A-C). Immunocytochemistry of positively sorted CD146 cells confirmed expression of CD146 in the majority of cells (Fig. 5D). Comparison of cell numbers pre and post MACS showed that approximately 2% of unsorted cells were CD146 positive (Fig. 5E), providing further emphasis on the rarity of CD146 cell subpopulations and requirements for optimal enrichment procedures. However, some CD146 positive cells were detected in negative fractions (approx. 14%) (Fig. 5F).

**Figure 5.**
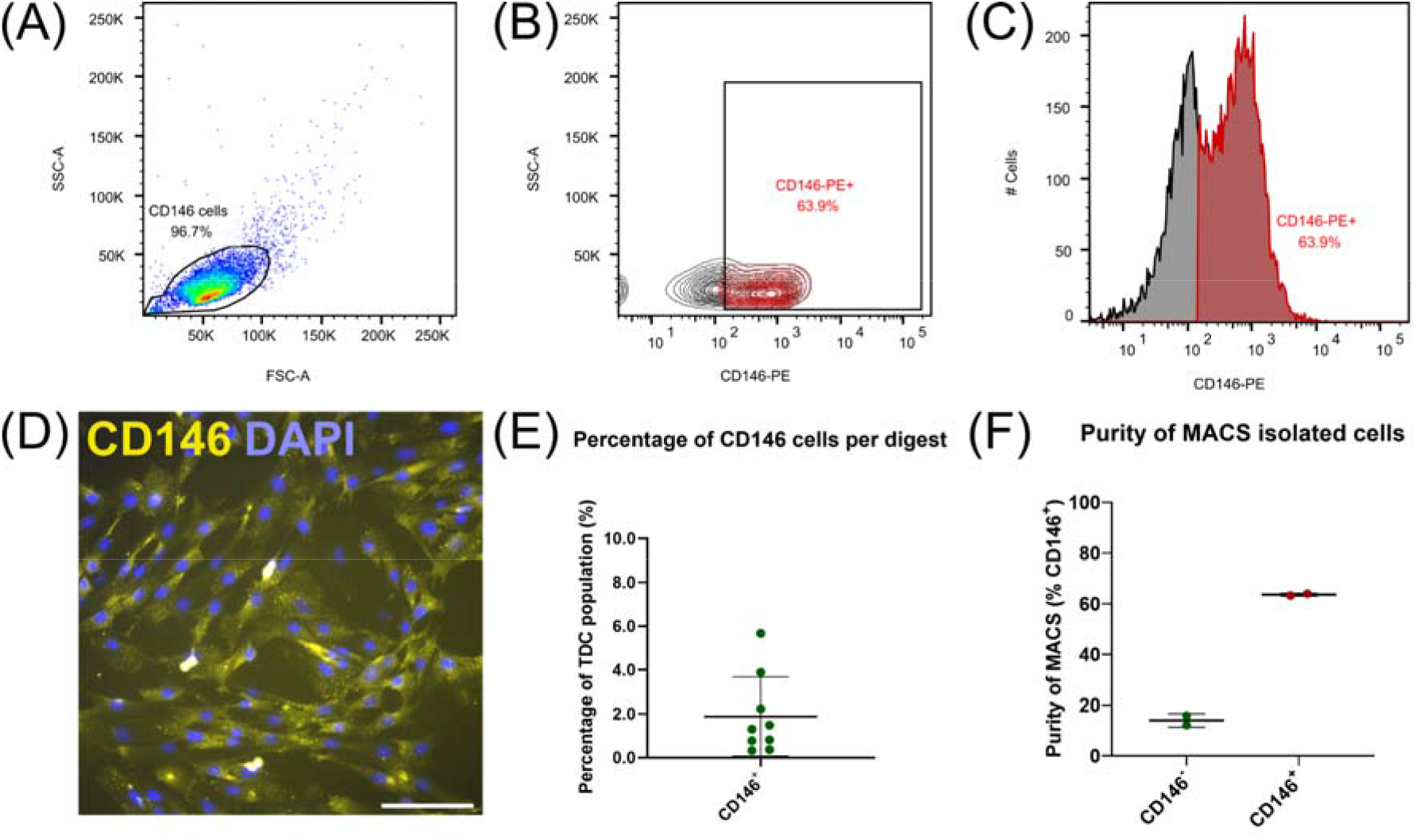
MACS yields good purity CD146^+^ cell isolations which can be expanded *in vitro* for downstream analyses. Flow cytometry (A-C) confirmed CD146 expression in CD146^+^ cell populations. Total events = 10,000. Immunocytochemically labelled CD146 cells (D; passage 2) labelled with CD146 reaffirmed that expression persisted once expanded *in vitro*. n=3 (biological replicates). Scale bar = 100 μm. (E) The percentage of CD146 cells (approx. 2%) yields by MACS from the original tendon-derived cell suspensions as determined by cell counting, reiterated the rarity of CD146 cells. n=9 (biological replicates). Across MACS isolations (F), flow cytometry confirms CD146 purity, albeit approximately 15% of cells in the CD146^-^ negative fraction were positive for CD146. n=2 (biological replicates) per cell fraction. Graphs were plotted as mean (μ) ± SD.

### 3.4 Interfascicular CD146 cells have limited differentiation potential

To assess their clonogenicity and multi-lineage potential, unsorted TDCs, CD146^+^, CD146^-^ cells were subjected to clonogenic, osteogenic and adipogenic assays using MSCs as a positive control. CD146^+^ cells showed no enhanced clonogenicity compared to CD146-negative cells or heterogenous TDCs (Fig. 6A-D). For adipogenesis, TDCs, CD146^+^ and CD146^-^ cells all showed limited adipogenic potential when stimulated (Fig. 6E-H). Under osteogenic conditions, unsorted TDCs displayed extensive calcium deposition with some mineralised nodules present, however virtually no calcium deposition nor mineralisation was detected in either CD146^+^ and CD146^-^ sorted cell populations (Fig. 6M-P).

**Figure 6.**
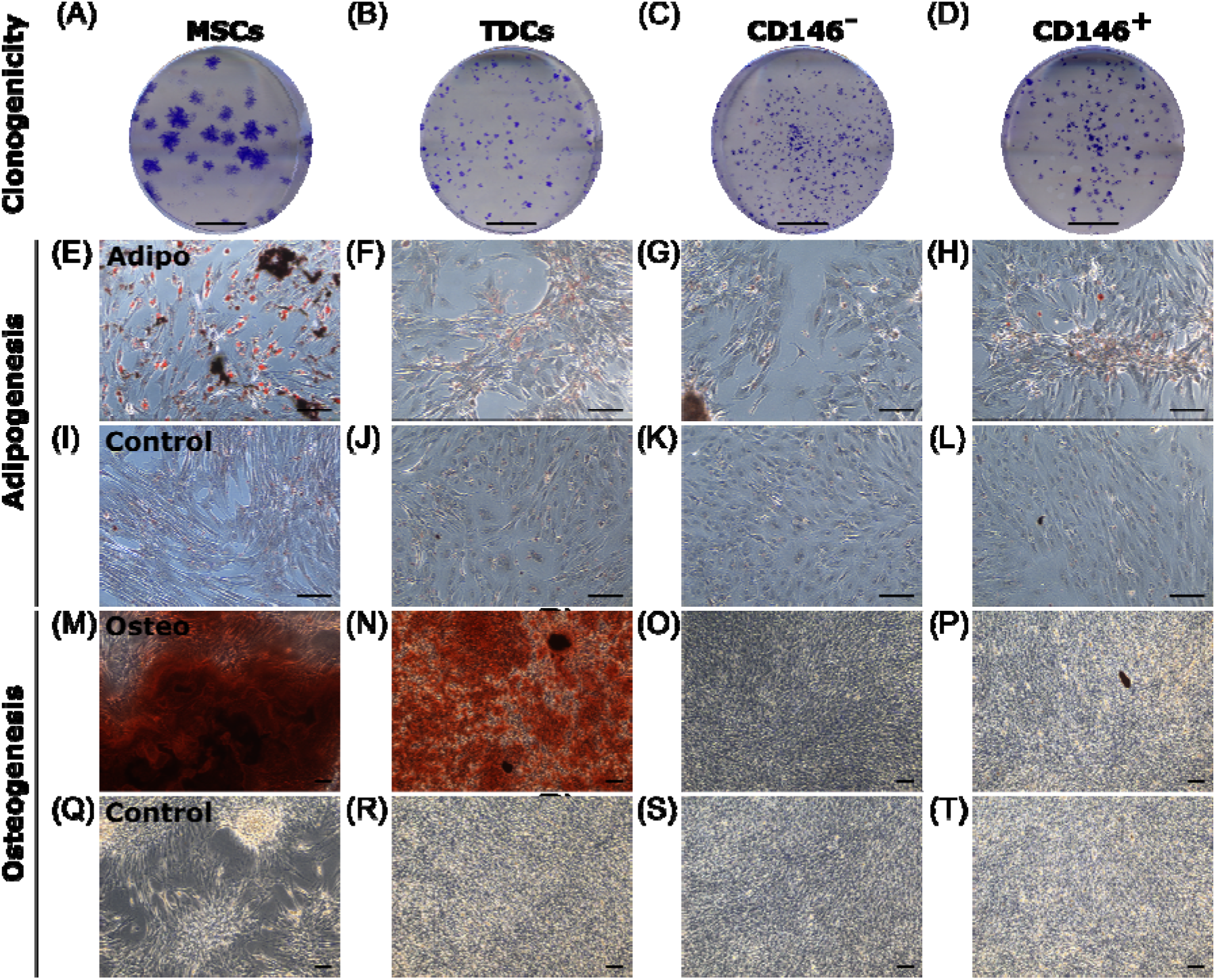
CD146^+^ cells exhibit limited clonogenicity and lineage potential. Representative images of colonies formed by MSCs, TDCs, CD146^-^ and CD146^+^ populations (A-D). Scale bar = 1 cm. Oil Red O staining of MSCs, TDCs, CD146^-^ and CD146^+^ cells (E-L) under adipogenic conditions using StemPro® Adipogenesis differentiation media (E-H) and control conditions (I-L) demonstrate that TDCs, CD146^+^ and CD146^-^ cells produce a limited number of lipid vesicles. Lipid vesicles = red. n=3 per cell type (biological replicates). n=3 per condition (technical replicates). Scale bar = 100 μm. Alizarin Red S staining of MSCs, TDCs, CD146^-^ and CD146^+^ cells (M-T) under osteogenic conditions containing 2 mM DiP (M-P) and control conditions (Q-T) demonstrate that tendon cells exhibit limited mineralisation capacity when separated. Mineralised nodules = black. Calcium deposits = red. Unmineralised matrix = reflective/white. n=3 for each cell type (biological replicates). n=2-3 wells for each condition (technical replicates). Scale bar = 100 μm.

## 4. Discussion

In this study, we have characterised CD146^+^ cell populations and their niche within the tendon IFM. We demonstrate that CD146^+^ cells exclusively localise to the IFM in healthy tendon and reside in a niche containing vascular basement membrane and vascular-associated proteins. In contrast to our hypothesis that the IFM is a progenitor cell niche, CD146^+^ cells exhibited limited differentiation potential, indicating they are unlikely to be stem/progenitor cells, and may instead be of vascular origin.

Several studies have demonstrated the presence of CD146^+^ cells in tendon; with recent single-cell RNA sequencing of human tendon revealing three cell populations that express CD146; one of which was an endothelial population which co-expressed CD31 [45]. Furthermore, in single-cell analyses of mouse tendon, CD146^+^ tendon cells, identified as haematopoietic cells, represented around 9% of TDCs [46]. In other tissues such as bone, CD31 and CD146 expression can be used to delineate endosteal and vascular populations which remodel the haematopoietic niche [47,48]. Previous research from our group has highlighted CD31 as an IFM-localised vascular marker [33] and the colocalisation of CD146 with CD31 we report here suggests that CD146^+^ may be an IFM population of pericytes/endothelial cells. It is notable that not all CD146^+^ cells *in situ* were CD31^+^, suggesting that CD146 labels more than one cell population within the IFM, or that CD31 expression may be transient.

In addition, our recent studies have established that CD146^+^ cells migrate to sites of injury in the rat Achilles tendon, which is accompanied by increased LAMA4 [30]. 3D imaging of tendon revealed an interconnected network of CD146 labelling within the IFM; structures which were similar those seen in 3D imaging of LAMA4 in SDFT [36]. The colocalisation of CD146 and LAMA4 in the current study further reinforces the putative ligand-receptor interaction that CD146 and LAMA4 share, which has been demonstrated in previous studies [49,50]. In chondrocytes, blockading LAMA4 inhibited cluster formation, which is typical of pathological cartilage, and also resulted in downregulation of Claudin-1 (previously identified as a tendon IFM protein) and MMP3 [51,52]. Recent studies have already established that loss of LAMA4 results in reduced CD146 cell expression and loss of basement membrane/niche maintenance in both mesenchymal and haematopoietic environments [53]. Therefore, LAMA4 may act as a homing receptor for migrating interfascicular CD146^+^ tenocytes, however the chemokines that facilitate this are yet to be identified.

Here, we confirm that both LAMA4 and LAMA5 are abundant within the IFM niche, alongside other vascular components ITGB1, VWF, EMCN, NTN1 and NRP1. During development, no other laminin chain functionally compensates for the α4-chain when knocked out during angiogenesis, however upon postnatal maturation, a compensatory upregulation of LAMA5 as a result of LAMA4 loss results in a milder vascular phenotype which suggests that the balance between laminin subunits LAMA4/LAMA5 ratios is critical for maintaining a healthy vascular network and vascular niche [54]. Given the abundance of LAMA4 and LAMA5, both chains and their full-length laminin isoforms 411 and 511 are likely essential to the IFM endothelial basement membrane due to their previously reported role in shear-stress response and mechanotransduction [55,56].

*In situ*, IFM cells were also positive for CD44 and CD90, which have been used as markers for putative stem/progenitor cell populations, both in tendon and other tissues, although their expression is likely acquired at later stages of differentiation and proliferation [57]. However, given that both markers were widely expressed throughout the IFM and fascicles, it is unlikely that these markers are specific for tendon stem cells in the equine SDFT and instead label several populations within tendon, including the tenocytes resident within fascicles. This is supported by single-cell RNA sequencing data from the mouse Achilles tendon showing that both CD44 and CD90 are expressed by tenocytes and other tendon cell populations [46].

The identification of multiple vascular structures using markers of endothelial/vascular cell lineages suggests the IFM houses a specialised vascular niche, rich in basement membrane proteins. This is supported by our previous proteomics data showing enrichment of basement membrane proteins in the IFM, including perlecan, laminins and collagen type IV [7]. The identification of perlecan-rich vascular networks in tendon IFM has major implications for the study of tendon. During development, perlecan is integral for tight packaging of interstitial tissues, which house vasculature, to ensure that maturation of endothelial tissues proceeds [58]. In addition, lymphangiogenesis within interstitial tissues is defined by the expression of perlecan and interstitial fluid flow [59]. In tendons, fascicular sliding may therefore be integral to IFM lymphatic and vascular remodelling. Moreover, VWF is likely to act as an endothelial cell ligand within the interfascicular basement membrane. It is notable that several of the network predicted CD146 interactions are basement membrane components, and localise to the IFM, indicating tethering of CD146 cells to the basement membrane.

*In vitro*, tendon derived cells showed similar protein expression to that seen *in situ*, with abundant labelling of CD44 and CD90, and limited labelling for CD146, which was expressed by 2% of cells derived from the SDFT. This is somewhat lower than the 9% of cells in the mouse Achilles that expressed CD146 as determined by single cell sequencing [46]; this discrepancy may be explained by species-specific differences. The equine model is a highly relevant and well-accepted model for tendon research as the SDFT and human Achilles share similar function, structure and injury risk [60,61]. However, we did observe some species-specific differences; for example, there was no detection of Nestin in equine TDCs, which is abundant in mice and human Achilles tendons, particularly in the IFM [21]. Another explanation for the discrepancy in population proportions in this study is the removal of the epitenon in the current study, which is known to house CD146^+^ cells [36]. MACS was successfully employed to enrich CD146 populations, with approximately 65% of cells positive for CD146 post-sorting as determined by flow cytometry. This percentage is likely an under-representation, as CD146^+^ cells were detected using flow cytometry immediately post MACS-enrichment, such that some CD146 antigens may still be bound to the magnetic label used during MACS, meaning they were not available for fluorescently tagged antibodies to bind to and so were not detected using flow cytometry. Indeed, immunocytochemistry of CD146^+^ cells showed that virtually all cells labelled positively for CD146 post-MACS enrichment. However, a proportion of negatively selected cells were CD146^+^ post-sorting, likely due to a small number of CD146^+^ cells not binding to the column and therefore being eluted with the negative fraction. Purity could have been improved by additional rounds of sorting; however, this would have resulted in insufficient cell numbers for downstream experiments.

All cell populations exhibited similar clonogenic and limited differentiation potential, which agrees with previous studies that demonstrated equine SDFT-derived TSPCs have limited differentiation potential [62]. Similarly, proliferation rates did not change significantly with cell types expanded under normoxic conditions, as reported in the same study. However, in the current study, lipid vesicles were produced in both adipogenic-induced TDCs and CD146^+^ cells; in contrast to the above study where adipogenesis was not detected in stimulated TSPCs [62]. Unsorted tendon cell populations appear to have greater osteogenic potential compared to sorted CD146 populations. The increased mineralisation in TDCs as opposed to both sorted populations suggests that osteogenic capacity is increased when crosstalk between CD146^+^ cells and CD146^-^ is possible in culture. Given that CD146 is described as a marker of pericytes, co-culturing CD146 vascular cells with tenocytes may enhance their calcification potential further, as reported in atherosclerotic tissues which described greater mineralisation in zones of CD146-expressing pericytes [63]. We were unable to assess chondrogenesis due to limited cell numbers of CD146^+^ cells after sorting, which were not sufficient for micromass survival during chondrogenic pellet induction. Taken together, CD146^+^ cells do not exhibit stem cell plasticity and are likely a population of pericyte-like cells. While the multipotency of pericytes has been demonstrated in a range of species [64], other studies have shown that pericyte plasticity varies between tissue types, with some pericytes having limited differentiation potential [65]. It is possible that, while tendon pericytes have a limited multipotency, they can differentiate down a tenogenic lineage, and indeed single cell sequencing data indicate that pericytes are a source of progenitor cells for adult tenocytes in murine tendon [66]. As tendon CD146^+^ populations have been shown to migrate to sites of injury, establishing further understanding of their local microenvironment, lineage origins, *in vitro* characteristics, and the effects of ageing will aid future research aimed at establishing if mobilising these populations can enhance intrinsic repair.

## 5. Conclusions

In tendon, CD146 demarcates an IFM-specific cell population that reside in a niche rich in basement membrane and vascular proteins. CD146^+^ cells have limited clonogenicity and differentiation potential indicating they are unlikely to be stem/progenitor cells. Instead, co-localisation of CD146 with the vascular cell marker CD31 indicates these cells may be pericytes. As previous studies have shown that CD146 cells migrate to sites of injury, establishing regenerative strategies that utilise endogenous tendon pericyte cell populations to promote intrinsic repair could act as a viable and effective method for improving healing responses and preventing tendon re-injury.

## Supplementary Materials

Table S1: Antibodies used for immunolabelling and their blocking conditions; Figure S1: Negative and isotype control labelling of SDFT tissues; Figure S2: Negative (secondary antibody) control staining for immunohistochemical labelling of SDFT; Figure S3: Workflow for the determination of IFM and fascicular boundaries in longitudinal SDFT sections.

## Data Availability Statement

Data is contained within the article or supplementary material.

## Author Contributions

Conceptualization, NM, CTT; methodology, NM, AAF, DW, CTT; investigation, NM, DEZ; writing—original draft preparation, NM, CTT; writing—review and editing, NM, DEZ, AAF, DW, CTT, JD, AAP; supervision, CTT, JD, AAP; project administration, CTT; funding acquisition, CTT. All authors have read and agreed to the published version of the manuscript.

## Institutional Review Board Statement

Collection of equine tendon was approved prior to commencement of the study by the Royal Veterinary College Ethics and Welfare Committee (URN-2016-1627b).

## Acknowledgments

The authors would like to thank Dr Isabel Orris and Dr Lucie Bourne for support and guidance on mineralization cultures.

