## Supplementary Materials for "The tendon interfascicular basement membrane provides a vascular niche for CD146^+^ pericyte cell subpopulations"

**Table S1**. Antibodies used for immunolabelling and their blocking conditions.

| Primary antibody | Supplier | Dilution | Secondary antibody (if applicable) | Supplier (if applicable) | Dilution (if applicable) | Blocking conditions |
| --- | --- | --- | --- | --- | --- | --- |
| CD31 | Abcam (ab28364) | 1:100 | Goat anti-mouse Alexa Fluor® 488 | Thermo Fisher Scientific (A-11001) | 1:500 | 5% goat & 5% horse serum; 1% BSA |
| CD44 | Bio-Rad (MCA2219GA) | 1:100 | Goat anti-mouse Alexa Fluor® 633 | Thermo Fisher Scientific (A-21052) | 1:500 | 5% goat & 5% horse serum; 1% BSA |
| CD90 (THY-1) | Fisher Scientific (15227117) | 1:100 | Goat anti-mouse Alexa Fluor® 633 | Thermo Fisher Scientific (A-21052) | 1:500 | 5% goat & 5% horse serum; 1% BSA |
| CD133 (PROM1) | St. John’s Laboratories (STJ96557) | 1:150 | EnVision peroxidase labelled polymer | Dako (K4065) | N/A  (Used neat) | 5% goat & 5% horse serum; 1% BSA |
| CD146 (MCAM) | Abcam (ab75769) | 1:100 | Goat anti-rabbit Alexa Fluor® 488; Goat anti-rabbit Alexa Fluor® 555; Goat anti-rabbit Alexa Fluor® 594 | Thermo Fisher Scientific (A- 11008); Abcam (ab150078); Thermo Fisher Scientific (A-11037) | 1:500 | 5% goat & 5% horse serum; 1% BSA |
| DAG1 | Millipore (05-593) | 1:200 | EnVision peroxidase labelled polymer | Dako (K4065) | N/A  (Used neat) | 5% goat & 5% horse serum; 1% BSA |
| EMCN | St John’s Laboratory (STJ92922) | 1:150 | EnVision peroxidase labelled polymer | Dako (K4065) | N/A  (Used neat) | 5% goat & 5% horse serum; 1% BSA |
| ITGB1 | St. John’s Laboratories (STJ93731) | 1:150 | EnVision peroxidase labelled polymer | Dako (K4065) | N/A  (Used neat) | 5% goat & 5% horse serum; 1% BSA |
| Laminin a4 (LAMA4) | St. John’s Laboratories (STJ93891) | 1:100 | Goat anti-rabbit Alexa Fluor® 594 | Thermo Fisher Scientific (A-11037) | 1:500 | 5% goat & 5% horse serum; 1% BSA |
| Laminin a5 (LAMA5) | St. John’s Laboratories (STJ93892) | 1:150 | EnVision peroxidase labelled polymer | Dako (K4065) | N/A  (Used neat) | 5% goat & 5% horse serum; 1% BSA |
| Mohawk homeobox (MKX) | Insight Bio (ARP32574_P050) | 1:100 | Goat anti-rabbit Alexa Fluor® 555 | Abcam (ab150078) | 1:500 | 5% goat & 5% horse serum; 1% BSA |
| Nestin (NES) | Abcam (ab22035) | 1:50 | Goat anti-mouse Alexa Fluor® 594 | Thermo Fisher Scientific (A-11032) | 1:500 | 5% goat & 5% horse serum; 1% BSA |
| Netrin-1 (NTN1) | St John’s Laboratory (STJ94407) | 1:150 | EnVision peroxidase labelled polymer | Dako (K4065) | N/A  (Used neat) | 5% goat & 5% horse serum; 1% BSA |
| Neuropilin-1 (NRP1) | St John’s Laboratory (STJ94438) | 1:150 | EnVision peroxidase labelled polymer | Dako (K4065) | N/A  (Used neat) | 5% goat & 5% horse serum; 1% BSA |
| Pan-laminin | Abcam (ab11575) | 1:500 | EnVision peroxidase labelled polymer | Dako (K4065) | N/A  (Used neat) | 5% goat & 5% horse serum; 1% BSA |
| Perlecan | Fisher Scientific (MA514641) | 1:500 | EnVision peroxidase labelled polymer | Dako (K4065) | N/A  (Used neat) | 5% goat & 5% horse serum; 1% BSA |
| Type IV Collagen | Southern Biotech (1340-01) | 1:200 | Rabbit anti-goat immunoglobulins/HRP | Dako (P0160) | 1:200 | 5% rabbit & 5% horse serum; 1% BSA |
| VWF | Dako (A0082) | 1:150 | EnVision peroxidase labelled polymer | Dako (K4065) | N/A  (Used neat) | 5% goat & 5% horse serum; 1% BSA |

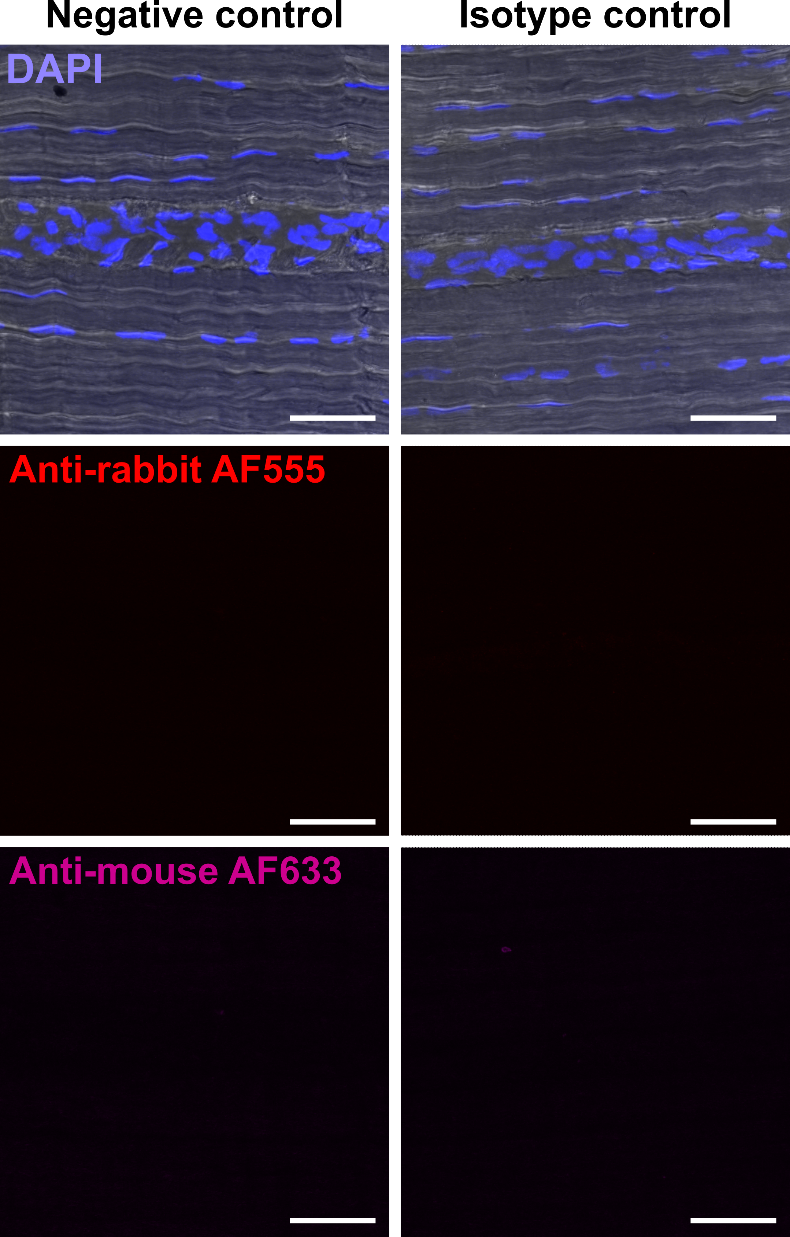

**Supplementary Figure S1.** Negative and isotype control labelling of SDFT tissues. Representative images of negative (left panel) control staining of SDFT tissue with goat anti-mouse Alexa Fluor® 633 and goat anti-rabbit IgG Alexa Fluor® 555 secondary antibodies applied only (1:100 for both). Isotype (right panel) matched control labelling was performed with rabbit IgG and mouse IgG isotype primary antibodies (1:100 for both) followed by incubation with secondary antibodies. Scale bar = 50 µm.

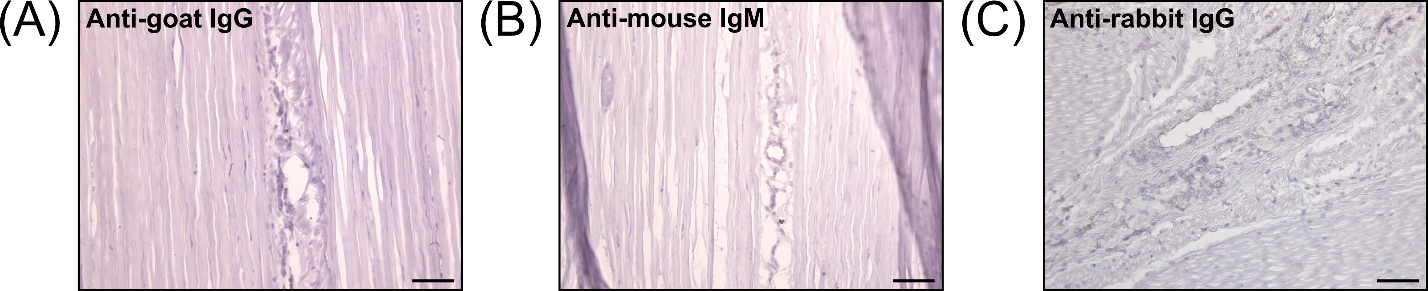

**Supplementary Figure S2.** Negative (secondary antibody) control staining for immunohistochemical labelling of SDFT. Longitudinal SDFT sections without primary antibodies labelled with (A) Dako rabbit anti-goat IgG secondary, and (B,C) EnVision peroxidase labelled polymer (conjugated to goat anti-mouse and goat anti-rabbit immunoglobulins). Scale bar = 75 µm.

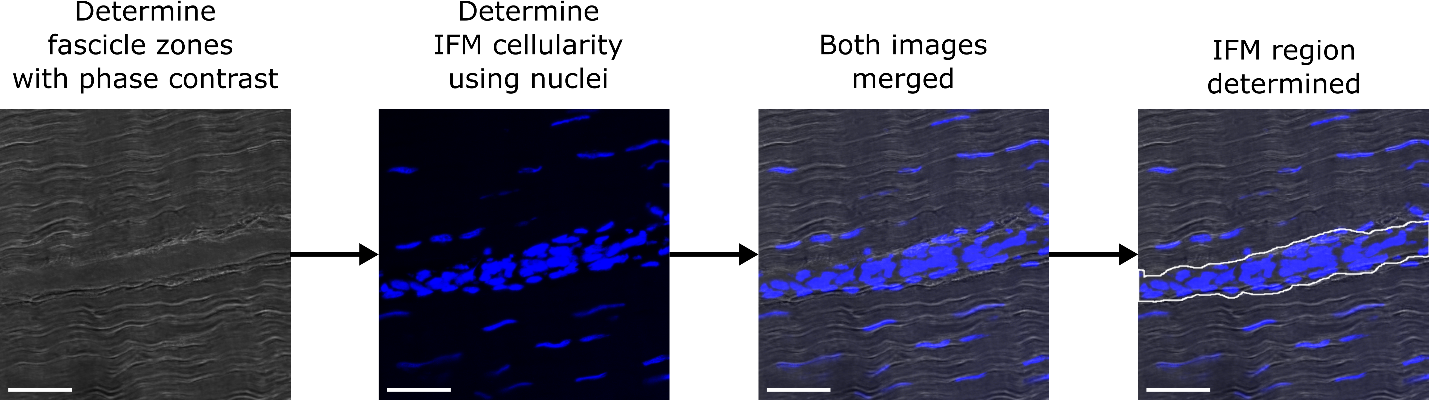
 **Supplementary Figure S3.** Workflow for the determination of IFM and fascicular boundaries in longitudinal SDFT sections. Phase contrast imaging (grey) and nuclei (DAPI = blue) aid in identifying IFM and fascicle boundaries (dashed line). Scale bar = 50 µm.
